# Axis convergence in *C. elegans* embryos

**DOI:** 10.1101/2023.03.27.534329

**Authors:** Archit Bhatnagar, Michael Nestler, Peter Groß, Mirna Kramar, Mark Leaver, Axel Voigt, Stephan W. Grill

## Abstract

Embryos develop in a surrounding that guides key aspects of their development. For example, the anteroposterior (AP) body axis is always aligned with the geometric long axis of the surrounding eggshell in fruit flies and worms. The mechanisms that ensure convergence of the AP axis with the long axis of the eggshell remain unresolved. We investigate axis convergence in early *C. elegans* development, where the nascent AP axis, when misaligned, actively re-aligns to converge with the long axis of the egg. Here, we identify two physical mechanisms that underlie axis convergence. First, bulk cytoplasmic flows, driven by actomyosin cortical flows, can directly reposition the AP axis. Second, active forces generated within the pseudocleavage furrow, a transient actomyosin structure similar to a contractile ring, can drive a mechanical re-orientation such that it becomes positioned perpendicular to the long axis of the egg. This in turn ensures AP axis convergence. Numerical simulations, together with experiments that either abolish the pseudocleavage furrow or change the shape of the egg, demonstrate that the pseudocleavage furrow-dependent mechanism is the key driver of axis convergence. We conclude that active force generation within the actomyosin cortical layer drives axis convergence in the early nematode.

---

The development of a single-cell embryo into an adult animal features the establishment of up to three anatomical body axes – anterior-posterior, dorsal-ventral and left-right. In general, these axes are intrinsic to the embryo (Goldstein and Freeman, 1997), yet consistently orient relative to geometric cues. In the nematode worm *C. elegans*, the anterior-posterior (AP) axis coincides with the geometric long axis of the ellipsoidal egg (Goldstein and Hird, 1996). However, during the establishment of the AP axis, the two axes may not always coincide. In this case, the AP axis actively converges towards the long axis of the egg (Goldstein and Hird, 1996). In this study, we seek to understand how the ellipsoidal geometry of the *C. elegans* egg directs this convergence of the AP axis with the long axis.

The orientation of the *C. elegans* AP axis is defined by the formation of mutually exclusive domains of PAR (partitioning-defective) proteins in the actomyosin cortex in the one-cell stage embryo, in particular the anterior (consisting of anterior PAR proteins, or aPARs) and posterior (consisting of posterior PAR proteins, or pPARs) domains (Guo and Kemphues, 1995; Rose and Gönczy, 2014; Motegi and Seydoux, 2013; Hoege and Hyman, 2013; Lang and Munro, 2017). These domains form as a result of cell polarity establishment about 30 minutes after fertilization. The egg is initially unpolarised, with aPARs uniformly enriched throughout the entire cortex (Cuenca et al., 2003; Cowan and Hyman, 2004; Schonegg and Hyman, 2006). Normally, centrosomes carried by the sperm-donated male pronucleus initiate polarity establishment at the site of sperm entry (O’Connell et al., 2000; Wallenfang and Seydoux, 2000; Cowan and Hyman, 2004; De Henau et al., 2020; Klinkert et al., 2019). Polarization proceeds via guided self-organized mechanochemical feedback between the PAR network (Motegi and Seydoux, 2013) and surface stresses and flows in the actomyosin cortex (Bois et al., 2011; Gross et al., 2017; Nitschke et al., 2019; Nestler and Voigt, 2022): Centrosomal microtubules load cytoplasmic pPARs onto the cortex (Wallenfang and Seydoux, 2000; Cowan and Hyman, 2004; Gross et al., 2019). Posterior PARs exclude myosin from the cortex, establishing myosin-poor domains (Motegi and Seydoux, 2013; Hoege and Hyman, 2013; Lang and Munro, 2017; Gross et al., 2019); the myosin imbalance between the pPAR domains and aPAR-rich regions generate active tension imbalances within the actomyosin cortex which drive cortical flow (Mayer et al., 2010; Bois et al., 2011). Cortical flows circulate away from the male pronucleus and towards the future anterior end (Motegi and Seydoux, 2013; Hoege and Hyman, 2013; Lang and Munro, 2017; Mayer et al., 2010), further redistributing PAR proteins by advective transport (Munro et al., 2004; Goehring et al., 2011; Mayer et al., 2010). Altogether, mutual antagonism between pPARs and aPARs and their transport by cortical flows leads to the formation of mutually exclusive domains of aPARs and pPARs: aPARs at the anterior and pPARs at the posterior (Gross et al., 2019); giving rise to a polarized cell and defining the embryonic AP body axis (Goldstein and Hird, 1996).

The ellipsoidal-like geometry of the *C. elegans* zygote is imposed by the eggshell surrounding the zygote, which forms shortly after fertilization (Johnston and Dennis, 2012). The zygote has one long axis about 50 µm in length, and two short axes about 30 µm in length (Begasse and Hyman, 2011; Riddle et al., 1997). How then is the AP axis oriented along the geometric long axis of the egg? As described above, the location of sperm entry imparts positional information to the cell polarity machinery. The sperm tends to enter the oocyte at the tip (Goldstein and Hird, 1996; Bienkowska and Cowan, 2012), possibly due to how the oocyte enters the spermatheca (Ward and Carrel, 1979; McCarter et al., 1999). In this case, the orientation of AP axis is along the long axis of the egg ellipsoid. However, occasionally the sperm enters away from the tip, yet the AP axis eventually converges with the geometric long axis (Goldstein and Hird, 1996). In these cases, the male pronucleus ‘posteriorizes’; that is, it moves toward the closest tip of the embryo (Kimura and Kimura, 2020; Goldstein and Hird, 1996). This dynamically reorients the AP axis towards the long axis of the embryo, until the two axes converge. However, the physical mechanisms that drive this process of axis convergence remain poorly understood.

How does the AP axis converge towards the long axis? Flows in the cytoplasm have been observed to drive repositioning of cellular structures in the cytoplasm (Goldstein and van de Meent, 2015; Quinlan, 2016; Deneke et al., 2019; Yi et al., 2011; Wang et al., 2020). Such cytoplasmic flows, as observed in the one-cell embryo (Niwayama et al., 2011), have previously been proposed to drive posteriorization (Goldstein and Hird, 1996; Bienkowska and Cowan, 2012), and thus AP axis convergence. These flows are created by anterior-directed cortical flows at the surface (Niwayama et al., 2011). Cytoplasmic flows, in turn, could transport the male pronucleus (Kimura and Kimura, 2020; Mittasch et al., 2018) towards the closest tip. We refer to this scenario as the cytoplasmic flow-dependent mechanism. Another possibility is that cortical flows drive axis convergence in *C. elegans* embryos via the pseudocleavage furrow (Nigon et al., 1960), a contractile ring-like structure that forms at the boundary between the aPAR and pPAR domains (Cuenca et al., 2003; Cowan and Hyman, 2004). The ring arises by compressive flow-driven alignment of actin filaments (Reymann et al., 2016). The pseudocleavage furrow is dispensable for establishing the AP axis (Rose et al., 1995), but might modulate the dynamics of AP axis formation (Aras et al., 2018). In particular, if the furrow does not lie in a plane orthogonal to the long axis of the egg, constriction of the furrow (Reymann et al., 2016) would reposition it to minimise its circumference, reorienting the furrow to be in a plane orthogonal to the long axis. We refer to this scenario as the pseudocleavage furrow-dependent mechanism.

To evaluate these two mechanisms, we measure cortical flows and quantify axis convergence during AP axis establishment. We develop a mathematical model of axis convergence that recapitulates the measured flows—both cortical and cytoplasmic. An experimental reduction of cortical flows demonstrate that they are required for axis convergence. Experiments and numerical simulations that diminish the pseudocleavage furrow demonstrate that the pseudocleavage furrow-dependent mechanism is the primary driver of axis convergence, but the cytoplasmic flow-dependent mechanism also contributes. Experiments and numerical simulations that change the shape of embryos demonstrate the contribution of geometry to the rate of axis convergence: rounder embryos exhibit slower axis convergence. This reduction in rate is consistent between numerical simulations and experimental observations. This effect can also be described by a minimal model focusing on the action of the pseudocleavage furrow. Taken together, our results demonstrate how actomyosin-dependent active surface mechanics and flows translate the ellipsoidal geometry of the embryo into the orientation of the AP axis.

## Results

### Cortical flows drive AP axis convergence

We first set out to understand how the AP axis converges with the long axis of the embryo. To this end we observed the behaviour of PAR domains during AP axis establishment, via time-lapse imaging of the one-cell *C. elegans* embryo during this phase. We used a *C. elegans* line expressing PAR-6::mCherry (marking the anterior PAR domain) together with PAR-2::GFP (marking the posterior PAR domain) to visualise the PAR domains (see SI: Table 1). In agreement with previous observations (Goldstein and Hird, 1996), we found that in 33 out of 57 embryos where the AP axis is established off-axis (with a measured AP axis that started more than 10 deg out of convergence with the geometric long axis), the PAR domains moved such that within 7 minutes the AP axis converged with the long axis of the embryo (Figure 1a). We also find that AP axis convergence occurs concomitantly with the posteriorization of the male pronucleus (Figure 1b,c, male pronucleus visualised as a dark circle in the cytoplasm, see SI: Figure 1).

**Figure 1.**
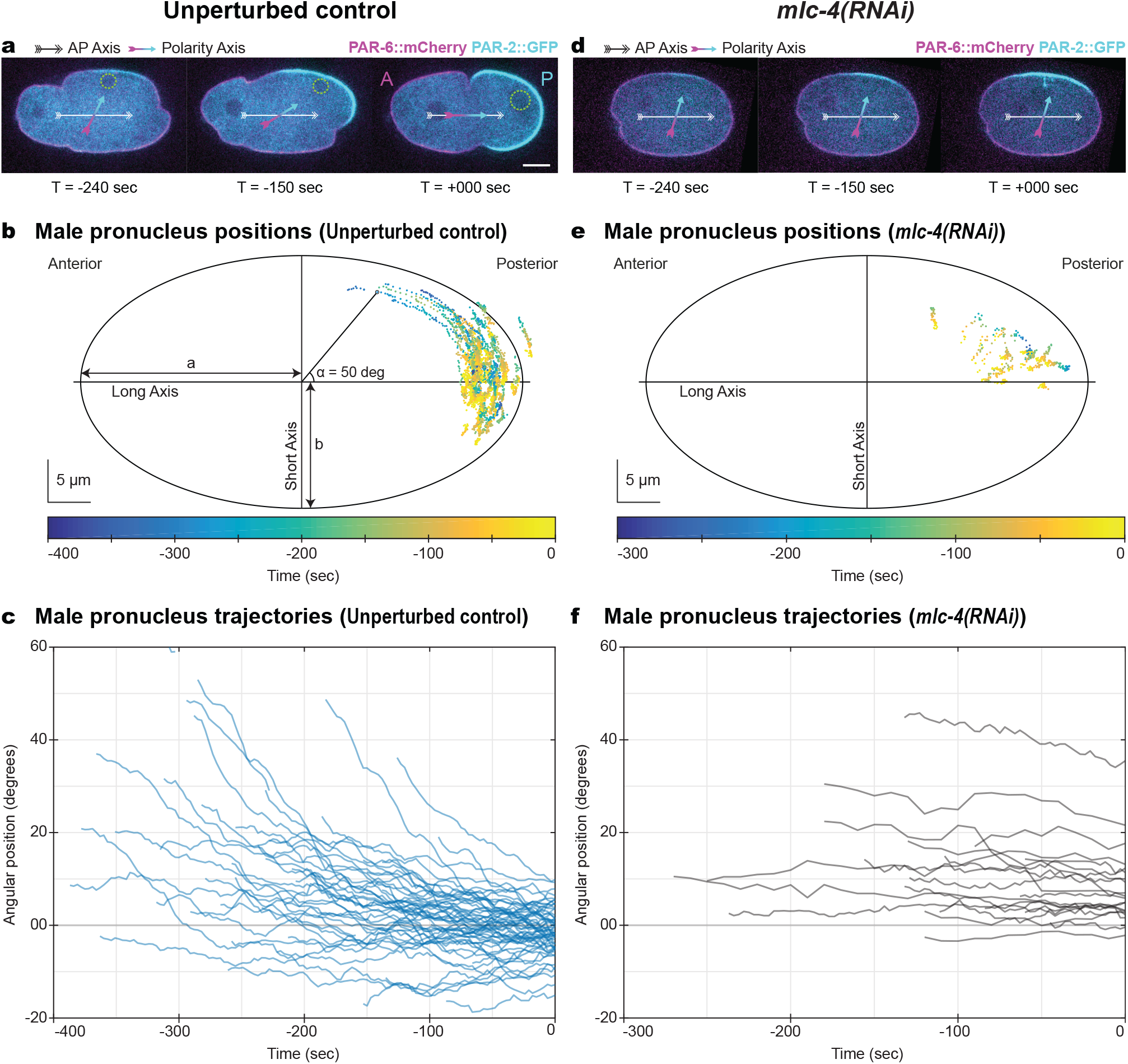
AP axis convergence and male pronucleus posteriorization depends on myosin activity. a) AP axis convergence in unperturbed control embryos is visualized by labeling anterior PAR (PAR-6::mCherry, in magenta) and posterior PAR (PAR-2::GFP, in cyan) domain. The male pronucleus (green dashed circle) moves with the posterior PAR domain, aligning the AP axis (cyan to magenta arrow) to the geometric long axis (White arrow). *T* =0 s is set at the end of posteriorization. Scale bar: 10 µm (for a,d). b) Male pronucleus positions (each point represents the center) during posteriorization in NMY-2::GFP labelled (Figure 1a) unperturbed embryos, color coded by time relative to end of posteriorization (see colorbar). Scale bar: 5 µm, Ellipse has semi-major axis *a* =28.4 µm and semi-minor axis *b* =17.5 µm lengths (for b,e). c) Angular position of the male pronucleus (on y-axis) as a function of time relative to end of posteriorization (on x-axis), in NMY-2::GFP labelled (Figure 1a) unperturbed embryos. Each line represents an individual embryo. d) No AP axis convergence is observed in *mlc-4* RNAi embryos, noted by no re-orientation of the posterior (cyan) and anterior (magneta) PAR domains. See (a) for details. e) Male pronucleus positions in *mlc-4* RNAi embryos labelled with NMY-2::GFP (Figure 1b) or PAR-6::mCherry and PAR-2::GFP, indicating lack of posteriorization in *mlc-4* RNAi embryos. See (b) for details. f) Angular position of the male pronucleus (on y-axis) as a function of time relative to end of posteriorization (on x-axis), in *mlc-4* RNAi embryos labelled with NMY-2::GFP (Figure 1b) or PAR-6::mCherry and PAR-2::GFP. Each line represents an individual embryo.

Furthermore, the angular velocity of the male pronucleus is highest when the angular position of the male pronucleus is largest; that is, the further the male pronucleus is from the posterior tip, the faster it moves toward the tip. Here, angular position refers to the angle between the long axis of the embryo and the line connecting the center of the male pronucleus to the center of the embryo. Specifically, we measured mean angular velocities of −2.1 ± 8.3 deg/s for angles between 0–10 deg compared to −18.9 ± 12.9 deg/s for angular positions between 30–40 deg (Figure 1c, negative velocities denote movements towards the posterior tip). Thus, for the range of angular positions measured (below approximately 45 deg), the rate of axis convergence is greatest when the axis is furthest displaced from the long axis.

Given the importance of cortical flows for AP axis establishment (Munro et al., 2004; Gross et al., 2019; Goehring et al., 2011), we next investigate their role in axis convergence. We used RNAi to impair function of *mlc-4*, an activator of myosin activity that regulates cortical flows (Shelton et al., 1999). In *mlc-4* RNAi embryos (Figure 1d, SI: Figure 1 b), flows were indeed reduced (see SI: Figure 3, SI: Figure 4 b,c, SI: Figure 5 b,c for comparison), the rate of axis convergence greatly diminished, and the AP axis failed to converge to the long axis (Figure 1d). Specifically, we measured mean angular velocities of −0.02 ± 0.04 deg/s for angular positions between 0–10 deg, −0.01 ± 0.03 deg/s for angular positions between 30–40 deg, with 10/18 embryos failing to re-orient their PAR domains (Figure 1e,f). Because MLC-4 activates myosin, and myosin is required for cortical flows (Shelton et al., 1999), a straightforward interpretation is that cortical flows are required for AP axis convergence (Zonies et al., 2010; Motegi et al., 2011; Tse et al., 2012).

### A mathematical model accounts for AP axis convergence

To understand how cortical flows could drive axis convergence, we developed a mathematical model. The model considers both active surface flows in the cortex itself (Mayer et al., 2010) and bulk cytoplasmic flows that are a consequence of cortical flows (Niwayama et al., 2011) (see Figure 2 and SI: section 2). This mathematical model should allow us to answer two questions: 1) Do both mechanisms, the cytoplasmic flow dependent mechanism and the pseudocleavage furrow-dependent mechanism (Figure 2a), contribute to axis convergence and, if so, to what extent? 2) Are the two mechanisms together sufficient to capture the spatiotemporal dynamics of the entire process?

**Figure 2.**
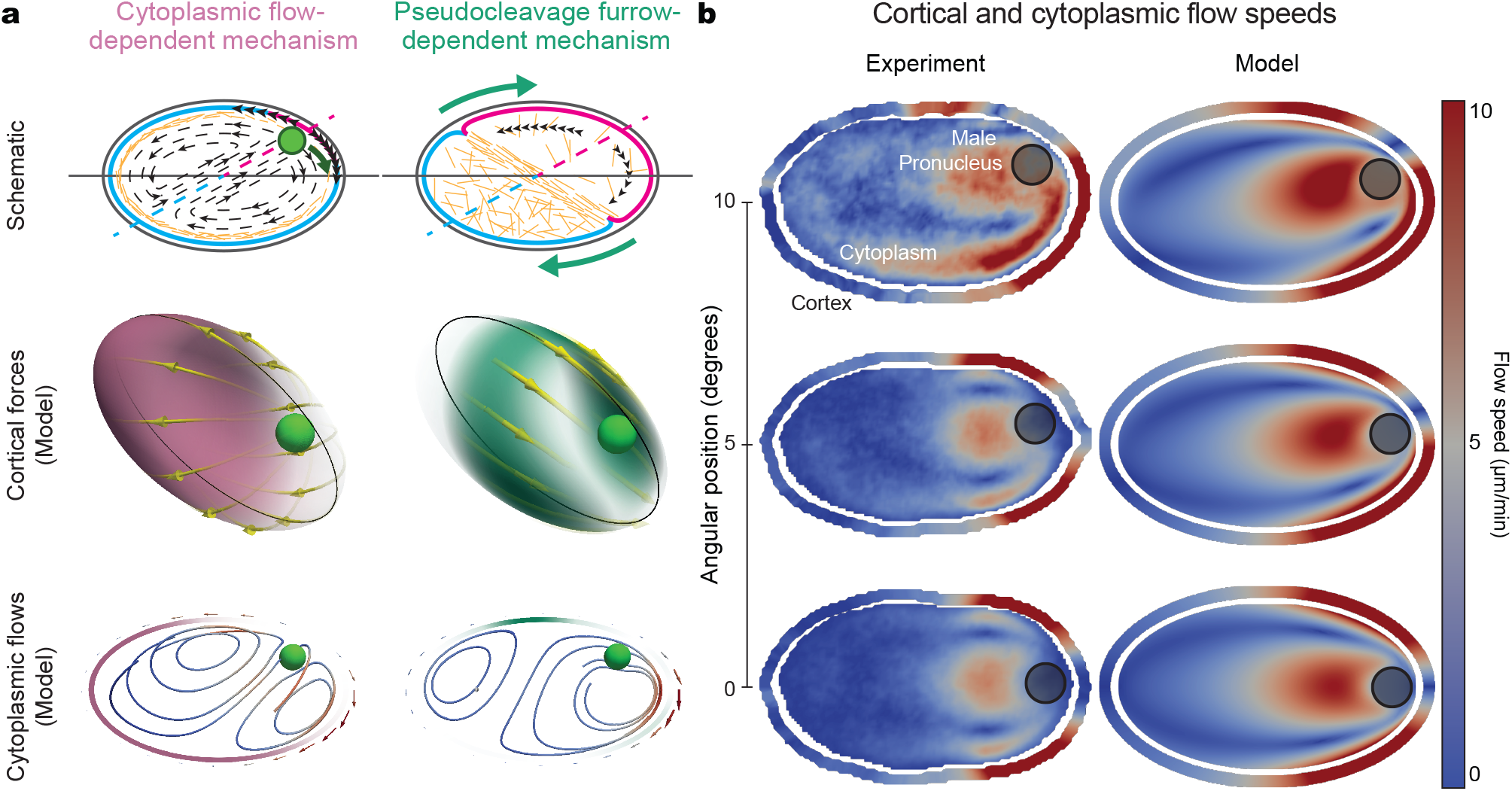
Numerical simulations capture the experimentally observed cortical and cytoplasmic flows during AP axis establishment in unperturbed embryos. a) Schematic detailing the two proposed mechanisms - cytoplasmic flow-dependent mechanism (left) and pseudocleavage furrow-dependent mechanism (right). 3D schematic depict the cortical forces induced by off-axis location of male pronucleus (green filled sphere) in the cortex: color represents magnitude (Transparent to red, increasing magnitude), green arrows represent direction. Black line indicates the 2D schematic boundary. In the 2D schematic, green circle represents the male pronucleus, outline around the embryo represents the cortex (transparent to red - increasing force magnitude), with black arrows representing cortical flow fields (magnitude - length of arrow). Streamlines inside the embryo represent cytoplasmic flows (blue to red - increasing speed). Each schematic represents the forces and flows that result from the corresponding mechanism alone - the full description considers both together, combined linearly. b) Cortical and cytoplasmic flow speeds, experimentally observed (left) compared to numerical simulations (right), for different male pronucleus positions (dashed green circle) in unperturbed embryos. Numerical simulations consider both cytoplasmic flow-dependent mechanism and pseudocleavage furrow-dependent mechanisms - see SI: Table 3, and SI: Figure 6. In each panel, ellipse interior represents cytoplasmic flows, and outer ellipse represents cortical flows. See colorbar for cortical flow velocities.

In the mathematical model (see SI: section 2), we approximate the geometry of the *C. elegans* embryo by a fixed ellipsoidal shape. The cortex at the surface of the ellipsoid is described in the general framework of active matter theory (Prost et al., 2015; Salbreux et al., 2012; Kruse et al., 2004) as an active nematic fluid layer (Kruse et al., 2005; Salbreux et al., 2009; Jülicher et al., 2018; Nestler and Voigt, 2022; Salbreux et al., 2022). Active stresses in this surface fluid layer arise through myosin activity (Mayer et al., 2010; Gross et al., 2019; Salbreux et al., 2012) and generate cortical flows. The cytoplasm itself is considered as an incompressible Newtonian fluid, with flows in it approximated as Stokes flow. At the interface between the cortex and the cytoplasm, the cytoplasm flows at the same velocity as the cortex; that is, cortical flows at the surface set a no-slip boundary condition for the bulk cytoplasm within the ellipsoid (Niwayama et al., 2011). The effect of bulk flows onto the surface fluid is captured by an effective frictional coefficient *γ* in the surface fluid layer (Mayer et al., 2010; Salbreux et al., 2012) (see SI: Equation 14). We assume that the friction between the embryo and the eggshell is negligible, and therefore consider only one friction term. With this coupling of surface and bulk flows, the cytoplasmic flow-field in the bulk is fully determined by the cortical flow-field on the surface (Niwayama et al., 2011) (see SI: Equation 15). The model considers the male pronucleus to be the organizer of active surface flows. The pronucleus is advected by the bulk cytoplasmic flows as a fluid particle with a large viscosity (Tanaka and Araki, 2000) (see SI: subsubsection 2.1.2). In addition, the male pronucleus is both in direct physical contact with the cortical layer and also attached to it via microtubules that radiate from the two centrosomes, which are adjacent to the male pronucleus, towards the cell cortex (Wallenfang and Seydoux, 2000; Kimura and Kimura, 2020; Bienkowska and Cowan, 2012). We capture the impact of these interactions by a drag coefficient between the male pronucleus and the cortical layer (*d*, see SI: Equation 18 and SI: subsubsection 2.1.2). The position of the male pronucleus determines the instantaneous distribution of myosin in the cortex and also defines the instantaneous AP axis (see SI: section 2, SI: Equation 9). The resulting gradients of myosin are responsible for the active stress distribution in the cortical layer which in turn drive cortical flows on the surface. The resulting cytoplasmic flows and its effect on the male pronucleus captures the cytoplasmic flow dependent mechanisms of axis convergence. The pseudocleavage furrow-dependent mechanism arises due to flow-induced local shear in the cortical layer, which causes actin filaments to align (Reymann et al., 2016). Active stresses depend on the state of filament alignment. Aligned active stresses generate the pseudocleavage furrow and drive rotation of the whole cell surface (Reymann et al., 2016; Miyazaki et al., 2015) (see Figure 2a and SI: section 2, SI: Equation 12). Thus the effects of actin alignment are embodied in the pseudocleavage furrow-dependent mechanism. Consequently, the distinction between the two mechanisms is whether they take actin alignment into account (Figure 2a). Altogether, for a given position of the male pronucleus, the mathematical model calculates cortical flows, cytoplasmic flows, and flow-derived male pronucleus velocity.

We implemented the model using the adaptive finite element toolbox AMDiS (Vey and Voigt, 2007; Witkowski et al., 2015). We solved for cortical flows on the ellipsoid surface using a surface finite element method for tangential vector- and tensor-quantities (Nestler et al., 2019). We solved the Stokes problem for cytoplasmic flows in the bulk using standard finite element methods within a diffuse domain (Li et al., 2009). We solved the coupled surface (cortex) and bulk (cytoplasm) flows concurrently. We selected numerical parameters such that the numerical solutions are stable and independent of these choices (see SI: subsection 2.2 and SI: subsection 2.3). Thus, the calculation of cortical flows, cytoplasmic flows, and flow-derived pronucleus velocity depends only on the four model parameters: hydrodynamic length *λ*_*H*_, active force relaxation *λ*_*A*_, nematic stress relaxation *λ*_*N*_ and drag coefficient between the cortex and male pronucleus *d* (see SI: Equation 22, SI: Table 3, SI: subsubsection 2.1.2), together with the two axes lengths that describe the geometry of the egg: semi-major axis *a* and semi-minor axis *b*.

We sought model parameters that recapitulate the cortical and cytoplasmic flows we observed experimentally in unperturbed embryos. For this we set the axes of the embryo ellipsoid to match the average axes lengths (with semi-major axis length *a* =28.4 µm and semi-minor axis length *b* =16.4 µm, SI: Figure 11 a). Because cortical flows are the basis for cytoplasmic flows, we sought parameters that result in calculated cortical flows that best match the experimental cortical flows (see SI: subsection 2.2). Cortical flows arise from myosin depletion near the male pronucleus; based on previous work (Gross et al., 2019) the model depletes myosin within 15.9 µm of the male pronucleus. With the ellipsoid dimensions and depletion distance set, we numerically solved the model for cortical flows over a range of angular positions of the male pronucleus (0 deg to 19 deg; SI: Figure 6). Our calculated cortical flows best match our experimentally measured cortical flows with the following parameter values: hydrodynamic length *λ*_*H*_ = 10 µm, active force relaxation *λ*_*A*_ = 11.5 µm^2^ s^−1^ and nematic stress relaxation *λ*_*N*_ = 152.5 µm^2^ s^−1^ (SI: Table 3, see SI: subsection 2.2 for calibration details). Bulk cytoplasmic flows are determined uniquely from cortical flows via the no-slip boundary condition. Calculated cytoplasmic flows show good agreement with experimental cytoplasmic flows (Figure 2b). Thus, for an optimal set of model parameters, our model faithfully recapitulates experimental cortical and cytoplasmic flows observed in unperturbed embryos.

Given that the mathematical model captures surface and bulk flow, we next evaluated how well the model predicts the movement of the male pronucleus towards the posterior tip. Specifically, we quantified: 1) angular position, taken as the angle between the line connecting the male pronucleus and embryo center to the long axis (Figure 3b), and 2) posteriorization velocity, taken as the component of the male pronucleus velocity tangential to the cortex, at the point on the cortex closest to the pronucleus (Figure 3c). The remaining model parameter that is needed to calculate male pronucleus dynamics is the drag coefficient *d*, which reflects direct interactions between the male pronucleus and the cortex (SI: subsubsection 2.1.2). Using model parameters that give the best match to experiment for cortical and cytoplasmic flows, we determined that a value of *d* =0.61 gives the best match to experimentally observed posteriorization velocities (for angular positions between 0–20 deg, Figure 3c). By integrating these best-match velocities, we obtain a predicted trajectory of the male pronucleus that agrees well with our experimental observations (SI: subsubsection 2.3.2, Figure 3b). Because numerical simulations closely match experimental measurements, we conclude that our model faithfully recapitulates the dynamics of the male pronucleus.

**Figure 3.**
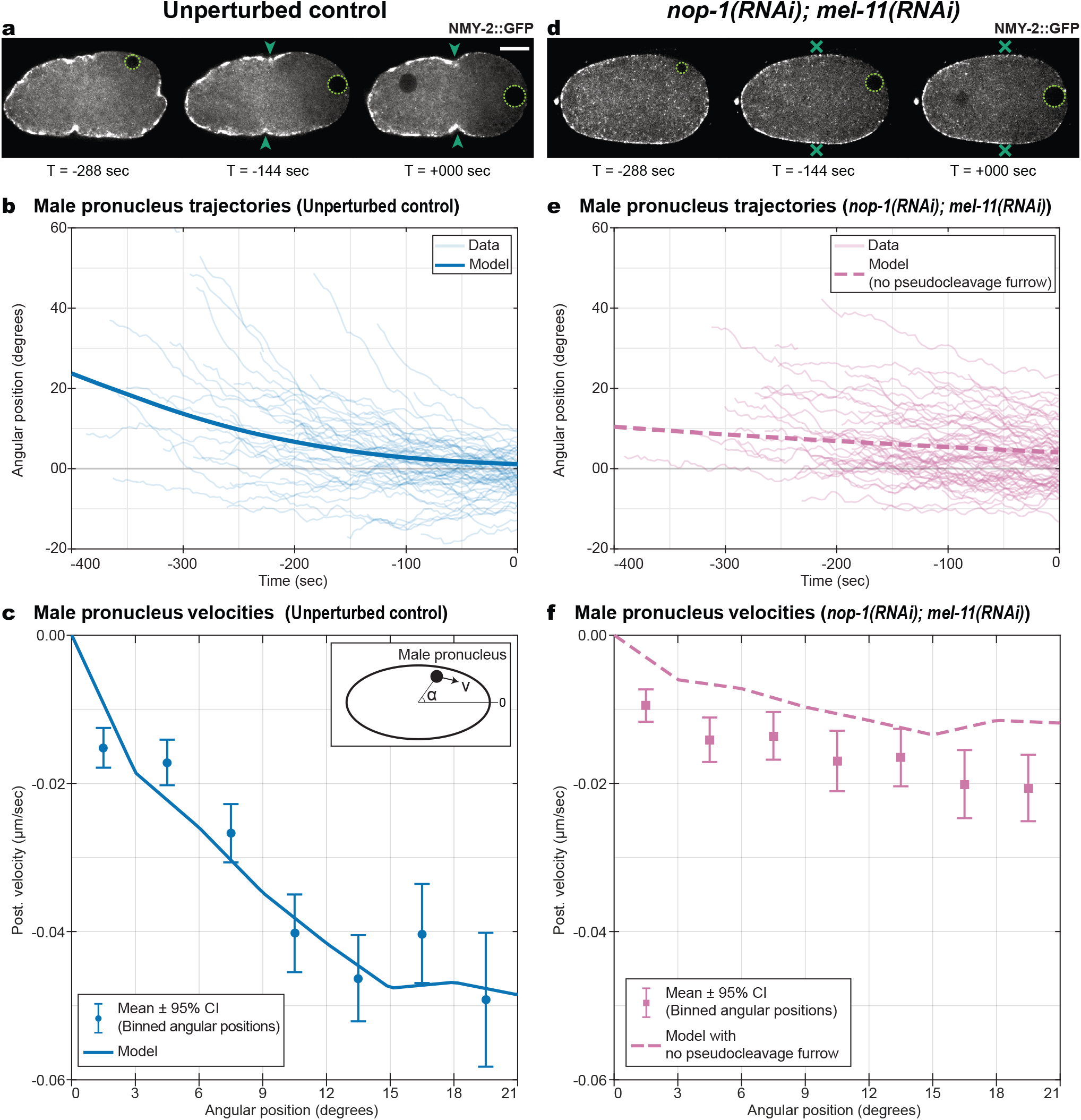
Experimental removal of pseudocleavage furrow via *nop-1;mel-11* RNAi reveals the role of pseudocleavage furrow in AP axis convergence. a,d) AP axis convergence observed in unperturbed control embryos (a) and *nop-1;mel-11* RNAi embryos (d) via time-lapse microscopy of embryos labelled with NMY-2::GFP (in gray). Arrows in (a) indicate the pseudocleavage furrow in unperturbed embryos, while crosses in (d) indicate the lack of the furrow in *nop-1;mel-11* RNAi embryos. *T* =0 s denotes end of posteriorization of the male pronucleus. Scale bar: 10 µm. b,e) Angular position of the male pronucleus (on y-axis, in deg), plotted as a function of time relative to end of posteriorization (on x-axis, in s) for unperturbed control embryos ((b), N = 57 - **same data as Figure 1c**) and *nop-1;mel-11* RNAi embryos ((e), N = 69). Blue thin lines in (b) and pink thin lines in (e) represent the experimentally observed trajectories of the male pronucleus in unpertured embryos (b) and *nop-1;mel-11* RNAi embryos (e) respectively; each line is an embryo. Blue thick line in (b) represents the predicted trajectory of the male pronucleus using the full mathematical model calibrated for unperturbed embryos. Pink dashed thick line in (e) represents the predicted trajectory using the mathematical model without the pseudocleavage furrow and calibrated for *nop-1;mel-11* RNAi embryos. (see SI: subsubsection 2.3.2 and SI: subsection 2.2) c,f) Posteriorization velocity of the male pronucleus (velocity component of the male pronucleus along the cell cortex, on y-axis, in µm/s) as a function of its angular position (on x-axis, in deg) in unperturbed embryos ((c), N = 57) and in *nop-1;mel-11* RNAi embryos ((f), N = 69). See inset in (c) for visual definition. Blue circles in (c) and pink squares in (f) denote the average posteriorization velocity for each angular position bin (bin size: 3 deg), with error bars representing the 95% confidence interval of the average, measured in unperturbed embryos (c) and *nop-1;mel-11* RNAi embryos (f) respectively. Blue curve in (c) and Pink dashed curve in (f) denotes the predicted posteriorization velocities using the full mathematical model (c) and mathematical model without pseudocleavage furrow (f) respectively (see SI: subsubsection 2.3.2 and SI: subsection 2.2 for details).

With this description in place, we are now in a position to evaluate to which degree the two mechanisms – the cytoplasmic flow-dependent mechanism and the pseudocleavage furrow-dependent mechanism – contribute to axis convergence.

### The predominant mechanism of AP axis convergence depends on the pseudocleavage furrow

In the mathematical model, the cytoplasmic flow-dependent mechanism and the pseudocleavage furrow-dependent mechanism are additive; furrow formation requires cortical flows, but the flows - although influenced by the furrow - do not require the furrow. Recall that the pseudocleavage furrow-dependent mechanism embodies the overall effect of actin alignment, of which the major effect is the formation of the furrow. Regardless, because the two mechanisms are separable, the model enables us to probe the function of the pseudocleavage furrow in AP axis convergence. Specifically, we can assess the importance of the pseudocleavage furrow dependent mechanism by modulating the *λ*_*N*_ parameter, which controls the actin alignment dependent active stresses. We observe that reducing the *λ*_*N*_ parameter leads to slower posteriorization velocities, but the AP axis still converges to the long axis (see SI: Figure 6). Thus, the model predicts that the pseudocleavage furrow-dependent mechanism is essential to achieve normal rates of AP axis convergence.

To verify this prediction experimentally, we generated embryos lacking a pseudocleavage furrow, but with normal cortical flows. To generate embryos with unaltered cortical flows, but without a pseudocleavage furrow, we performed double RNAi of *nop-1* and *mel-11*. NOP-1 modulates activity of the small GTPase RHO-1, a major regulator of actomyosin in the early embryo (Tse et al., 2012). Embryos from mothers that are mutant for NOP-1 lack a pseudocleavage furrow (Rose et al., 1995). MEL-11 is a myosin phosphatase (Piekny and Mains, 2002) that suppresses the activity of myosin in the cortex (Najafabadi et al., 2022). RNAi of *nop-1* eliminates the pseudocleavage furrow in the embryo (Tse et al., 2012), but cortical flows are also reduced (SI: Figure 3, SI: Figure 4 c, SI: Figure 4 d, SI: subsection 3.1). Under double *nop-1; mel-11* RNAi conditions, every embryo that we observed lacked a pseudocleavage furrow (69/69 embryos). Cortical flow speeds in the double-RNAi embryos were also comparable to unperturbed controls (average cortical flow speed: 3.34 ± 0.52 µm/min in double RNAi of *nop-1* and *mel-11*, as compared to 4.12 ± 0.59 µm/min for unperturbed control embryos, see SI: subsection 3.1, SI: Figure 3, SI: Figure 4 c, SI: Figure 4 d). Thus, *nop-1; mel-11* RNAi experimentally generates embryos lacking a pseudocleavage furrow, but with normal cortical flows.

To evaluate the prediction of the mathematical model, we measured the rate of convergence in *nop-1; mel-11* RNAi embryos. Qualitatively, the rate was slower (compare Figure 3e,f and Figure 3b,c). We calibrated our model based on the experimentally observed cortical flows in *nop-1; mel-11* RNAi embryos. Because we see no pseudocleavage furrow in these embryos, we set *λ*_*N*_ = 0 µm^2^ s^−1^, thereby excluding the pseudocleavage furrow dependent mechanism. Model parameters that yield the best match to experimental cortical flows for the *λ*_*N*_ = 0 µm^2^ s^−1^ condition are *λ*_*H*_ = 11 µm, *λ*_*A*_ = 7 µm^2^ s^−1^ (see SI: Table 3 and SI: Figure 7) which indicates that cortical mechanics are slightly altered in this condition. We leave the drag coefficient *d* =0.61 unchanged, and find that the model prediction for the *λ*_*N*_ =0 µm^2^ s^−1^ condition broadly matches both of the experimental readouts of AP axis convergence – posteriorization velocities and the male pronucleus trajectories – in *nop-1; mel-11* RNAi embryos (Figure 3e,f). Thus, the cytoplasmic flow-dependent mechanism accounts for the slow AP axis convergence in *nop-1; mel-11* perturbed embryos (Figure 3c,f; average posteriorization velocities, calculated over angular positions 0–20 deg: −0.01 ± 0.06 µm/s in *nop-1; mel-11* RNAi compared to −0.03 ± 0.06 µm/s for unperturbed embryos). We conclude that the predominant mechanism that drives AP axis convergence depends on the pseudocleavage furrow, as predicted by the mathematical model.

### Embryo geometry directs the rate of AP axis convergence

Embryo geometry is a prominent feature of the model, represented by the lengths of the long and the two equal short axes of the ellipsoid, *a* and *b* respectively. We focus on the aspect ratio *a*/*b* . To investigate how AP axis convergence depends on geometry, we calculated posteriorization velocities for embryos with different aspect ratios while holding the volume constant, setting all other model parameters to the values for unperturbed embryos (SI: subsection 2.2). Embryos with smaller aspect ratios show slower posteriorization velocities (SI: Figure 12 a, Figure 5a). Thus, the mathematical model predicts that rounder embryos have a slower rate of AP axis convergence.

To verify this prediction experimentally, we used RNAi to knockdown *ima-3* to generate embryos with aspect ratios smaller than unperturbed embryos (Sönnichsen et al., 2005). IMA-3 is a member of the importin *α* family of nuclear transport factors (Geles and Adam, 2001; Sönnichsen et al., 2005). Reducing *ima-3* levels by RNAi results in smaller, rounder embryos (Sönnichsen et al., 2005), (average *a/b* = 1.5 ± 0.2 for *ima-3* RNAi embryos compared to 1.8 ± 0.2 for unperturbed embryos, Figure 4a,b, SI: Figure 11 a). To obtain the model prediction for the rounder *ima-3* embryos, we used the same model parameters as the unperturbed embryos, but changed the aspect ratio of the ellipsoid to match the average *ima-3* RNAi embryos (SI: Table 3). Using the same model parameters is valid because both cortical flows and myosin concentrations in the cortex and cytoplasm are comparable between ima-3 RNAi and unperturbed embryos (see SI: subsection 3.2, SI: Figure 3, SI: Figure 4, SI: Figure 5 for cortical flows, SI: Figure 11 for myosin concentrations). The model prediction for this updated geometry matches both of the experimental readouts of AP axis convergence - male pronucleus trajectories and posteriorization velocities - in *ima-3* RNAi embryos (Figure 4c,d). Additionally, the experimentally-observed posteriorization velocities are faster in unperturbed embryos compared to *ima-3* embryos (Figure 4d; average posteriorization velocity over angular positions 0–20 deg: −0.03 ± 0.06 µm/s for unperturbed compared to −0.02 ± 0.06 µm/s in *ima-3* RNAi embryos). Consequently—as predicted by the model—rounder embryos have a slower rate of AP axis convergence.

**Figure 4.**
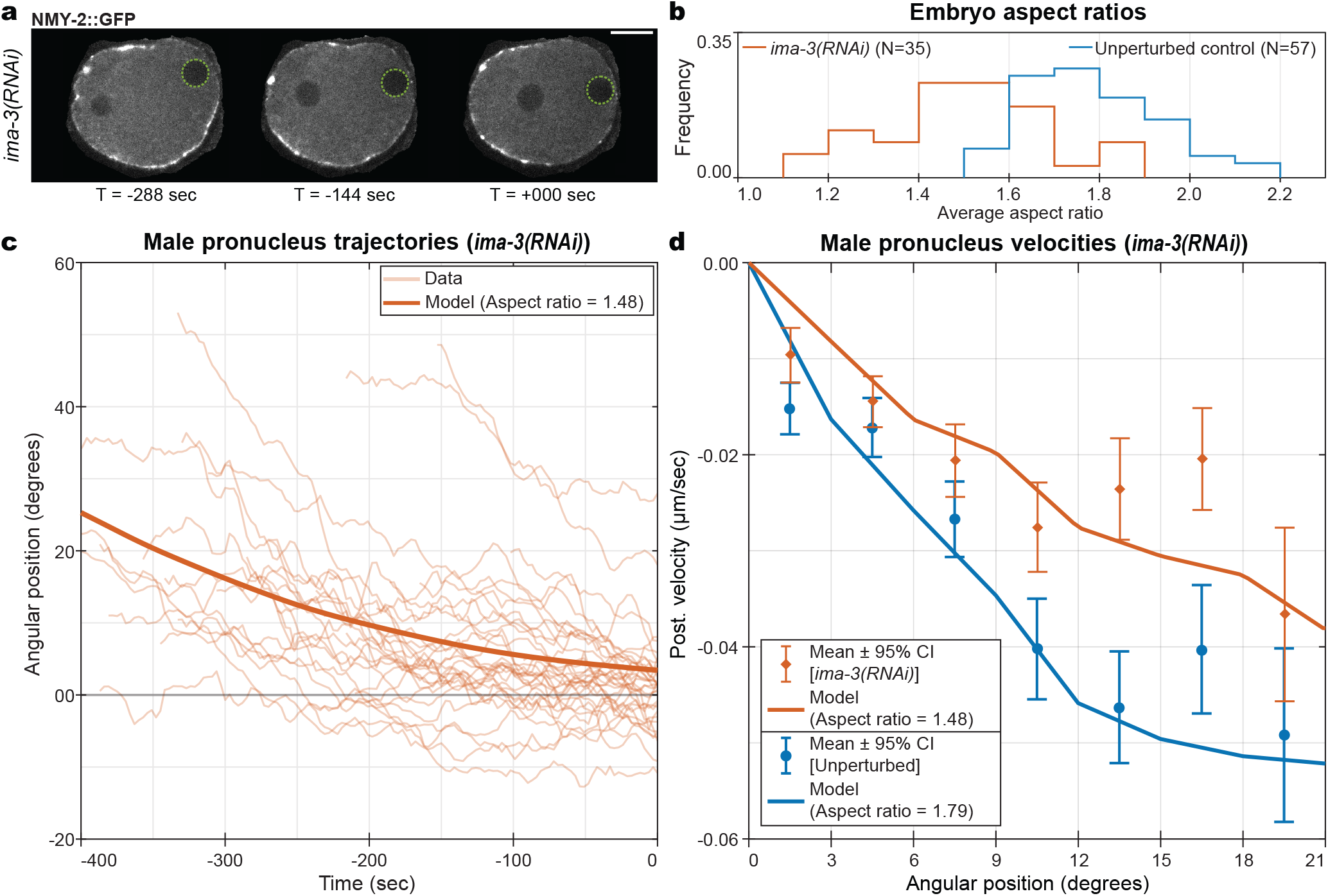
Rounder embryos generated by *ima-3* RNAi show slower AP axis convergence, consistent with predictions from mathematical model. a) AP axis convergence observed in rounder *ima-3* RNAi embryos via time-lapse microscopy of embryos labelled with NMY-2::GFP (in gray). *T* =0 s denotes end of posteriorization of the male pronucleus. Scale bar: 10 µm. b) Comparing aspect ratio (Length/Width) between unperturbed and *ima-3* RNAi embryos. Histogram depicts the distribution of average aspect ratio of an embryo in unperturbed embryos (Blue, N = 57, average aspect ratio = 1.8 ± 0.2) and *ima-3* RNAi embryos (Orange, N = 35, average aspect ratio = 1.5 ± 0.2) plotted as the frequency of embryos (fraction of embryos, along y-axis) observed in a given aspect ratio bin (along x-axis, bins of width 0.1). c) Angular position of the male pronucleus (on y-axis, in deg), plotted as a function of time relative to end of posteriorization (on x-axis, in s) for *ima-3* RNAi embryos. Orange thin lines represent the experimentally observed trajectories; each line is an embryo. Orange thick line represents the predicted trajectory of the male pronucleus using the full mathematical model, using the parameters from the unperturbed embryos (SI: Table 3) for unperturbed embryos (aspect ratio: 1.48). (see SI: subsubsection 2.3.2 and SI: subsection 2.2) d) Posteriorization velocity of the male pronucleus (on y-axis, in µm/s) as a function of its angular position (on x-axis, in deg) in unperturbed embryos (blue, N = 57, reproduced from Figure 3c) and in *ima-3* RNAi embryos (orange, N = 35). See inset in Figure 3c for visual definition. Blue circles and orange diamonds denote the average posteriorization velocity for each angular position bin (bin size: 3 deg), with error bars representing the 95% confidence interval of the average, measured in unperturbed embryos (blue circles) and *ima-3* RNAi embryos (orange diamonds) respectively. Curves denote the predicted posteriorization velocities using the full mathematical model, using the parameters from the unperturbed embryos (SI: Table 3) - blue for unperturbed embryos (aspect ratio: 1.79, reproduced from Figure 3b), orange for unperturbed embryos (aspect ratio: 1.48). (see SI: subsubsection 2.3.2 and SI: subsection 2.2 for details).

**Figure 5.**
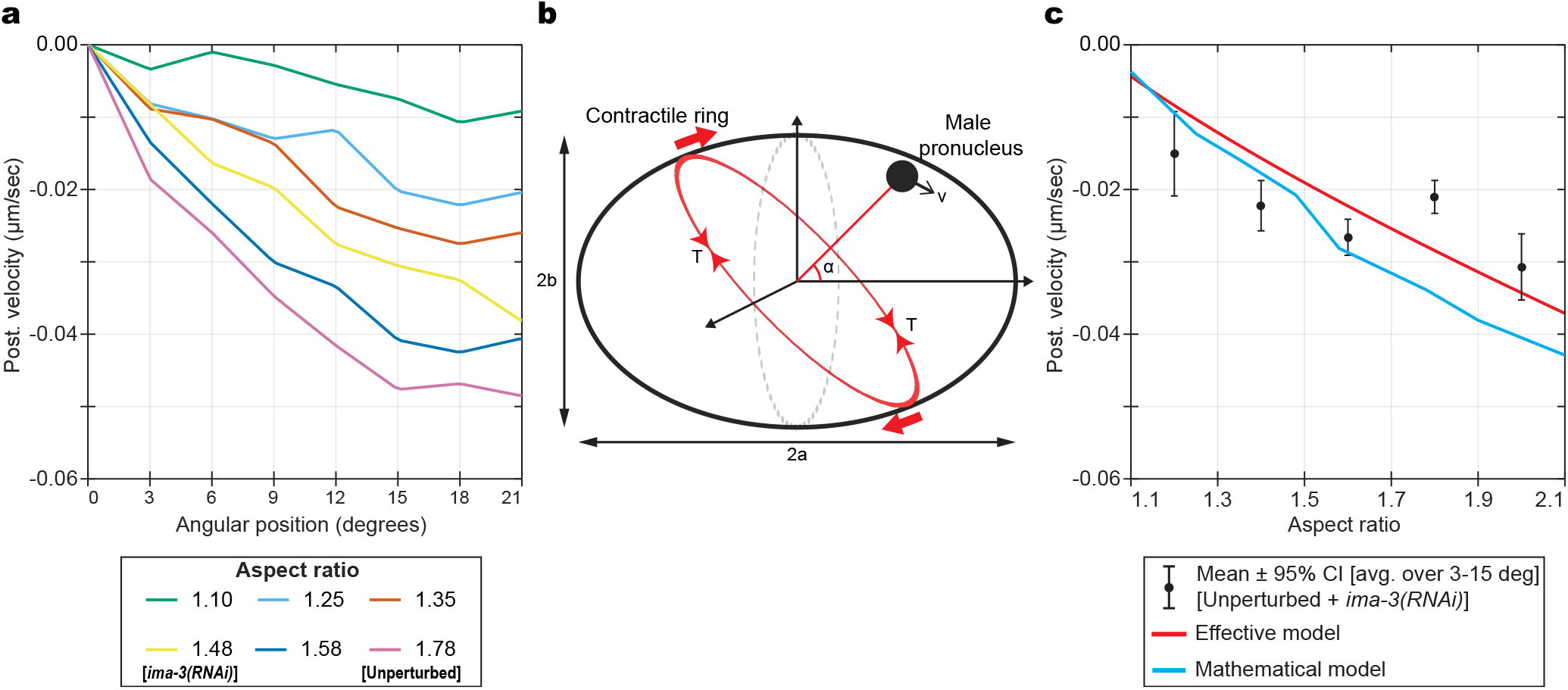
A minimal effective model can capture the relation between embryo geometry and rate of AP axis convergence. a) Predicted posteriorization velocity of the male pronucleus (on y-axis, in µm/s) as a function of its angular position (on x-axis, in deg) for embryo ellipsoid with different aspect ratios, using the full mathematical model. Only aspect ratio is varied, the volume of the ellipsoid is kept the same as the unperturbed embryos (SI: subsection 4.3). Aspect ratios 1.48 and 1.79 are used for predictions for *ima-3* RNAi (Figure 4d) and unperturbed embryos (Figure 3c,Figure 4d) respectively. b) Schematic depicting the effective model of a contractile ring (red ellipse) with fixed line tension (*T*) moving on an ellipsoid (black thick line, long axis of length 2*a* along x-axis and short axes of length 2*b* each along y- and z-axes). The ring is free to rotate around the z-axis (red block arrows). Its position is defined by the angle *α* between the long axis and normal to the plane of the ring, and is same as angular position of the male pronucleus. *v* is the posteriorization velocity of the male pronucleus. c) Posteriorization velocity of the male pronucleus (on y-axis, in µm/s) as a function of its aspect ratios (on x-axis). Black circles with error bars denote the observations from the combined unperturbed and *ima-3* RNAi embryos datasets, considering angular positions within 3–15 deg (N = 68). Black circles denote the average posteriorization velocity in each aspect ratio bin (bin size: 0.2). Error bars represent the 95% confidence interval in each bin. Curves denote the predictions from mathematical model (blue, SI: subsection 4.3) and effective model (red – 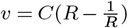, *R* = aspect ratio). For effective model, *C* =−2.32 × 10^−2^ µm/s (SI: section 4).

Because our mathematical model demonstrates a theoretical relation between embryo geometry and the rate of AP axis convergence – the more spherical the embryo, the slower the rate (SI: Figure 12 a, Figure 5a) – we sought to experimentally evaluate this relation directly. Instead of considering the rate of AP axis convergence – measured as the posteriorization velocity – as a function of angular position for a given average aspect ratio (Figure 4d), we now consider the posteriorization velocity as a function of aspect ratio. To obtain a characteristic velocity for each aspect ratio, we take the average of posteriorization velocity over an intermediate angular positions (between 3–15 deg, see Figure 5c). Given that cortical flows and myosin concentrations are comparable between unperturbed and *ima-3* RNAi embryos (see SI: subsection 3.2, SI: Figure 3, SI: Figure 4, SI: Figure 5 for cortical flows, SI: Figure 11 b,c for myosin concentrations), we combined the two datasets, thus considering aspect ratios between *a*/*b* = 1.1–2.1. We find good agreement between the model predictions and our experimental measurements (Figure 5c). Thus, we conclude that indeed: the more spherical the embryo, the slower the rate of AP axis convergence.

### A minimal model explains how geometry directs AP axis convergence

Why does the rate of AP axis convergence depend on embryo geometry? Our mathematical model and experimental results show that the predominant driver of AP axis convergence is the pseudocleavage furrow-dependent mechanism. In the mathematical model, the pseudocleavage furrow-dependent mechanism embodies the overall effect of actin convergence and associated active stresses, of which the major effect is the formation of the pseudocleavage furrow. In the embryo, the pseudocleavage furrow is a contractile ring. Thus, a simple, minimal model of AP axis convergence would be a contractile ring rotating on an ellipsoid. Is this minimal model sufficient to capture the relation between geometry and AP axis convergence?

In this minimal model (see SI: section 4), the ellipsoid has long axis of length 2*a* and short axis of length 2*b*. The centers of the ring and the ellipsoid are coincident. The angle between the normal to the plane of the ring and the long axis of the ellipsoid is *α*. The sole degree of freedom in the model is *α*; thus changes in *α* encapsulate all the dynamics of the model. Changes in *α* are opposed by a coefficient of friction (Figure 5b, SI: Figure 12 b) In addition, we assume that the male pronucleus moves with the same angular velocity as the ring. For small angles *α*, we obtain the following relation between the posteriorization velocity *v* of the male pronucleus and *α* (SI: section 4, SI: Equation 30):

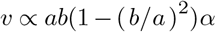

We find that to a reasonable extent, the simple effective model recapitulates both the experimentally observed and the simulated posteriorization velocities as a function of aspect ratio (Figure 5c). We conclude that this simple, minimal model captures the essence of the relation between geometry and AP axis convergence.

## Discussion

Here, we show that in *C. elegans* two partially redundant, additive mechanisms drive the convergence of the AP axis with the geometric long axis of the egg. Both are driven by flows in the actomyosin cortex, which behaves as an active nematic fluid layer. Flows in the cortical layer are organised by the centrosomes associated with the male pronucleus (Gross et al., 2019; Wallenfang and Seydoux, 2000). Cortical flows generate cytoplasmic flows and the pseudocleavage furrow. Both features translate the geometry of the ellipsoidal embryo to orient the AP axis towards the long axis of the embryo. Typically, both mechanisms are engaged, but the pseudocleavage furrow-dependent mechanism dominates.

Previous work has shown that the pseudocleavage furrow is largely dispensable for proper AP axis formation and orientation (Rose et al., 1995; Cowan and Hyman, 2004). We argue that in embryos without a pseudocleavage furrow, AP axis convergence takes place because the cytoplasmic flows advect the male pronucleus towards the closest tip. In embryos with a pseudocleavage furrow, AP axis convergence takes place rapidly, since the male pronucleus migrates quickly due to action of the furrow. Furthermore in pseudocleavage furrow-deficient embryos, incomplete AP axis convergence early in development could be corrected later by slow movement of PAR domains (Mittasch et al., 2018; Geßele et al., 2020). Such a correction may also be geometry-dependent, due to the tendency of PAR domains to form in regions of high curvature (Klinkert et al., 2019). Thus, multiple redundant mechanisms ensure proper positioning of the AP axis even if the pseudocleavage furrow fails to form.

To conclude, the AP axis converges to the long axis because cortical flows translate the geometry of the embryo into directed movement. The cortical layer is sensitive to geometry due to the nematic nature of the cortex: the alignment of actin filaments in the cortical layer depends on local curvature. Myosin motors acting on aligned filaments generate active anisotropic stresses physically embodied as the pseudocleavage furrow (Reymann et al., 2016). The nematic ordering of actin filaments thus gives rise to curvature-sensitive stresses that leads to the rotation of the pseudocleavage furrow, a directional movement. Previous studies have explored the relation between geometry and the behaviour of nematic materials (Giomi, 2015; Nitschke et al., 2018, 2020; Maroudas-Sacks et al., 2021; Nestler and Voigt, 2022; Hoffmann et al., 2022; Bell et al., 2022; Salbreux et al., 2022; Nitschke et al., 2022). Here, our mathematical model shows how the nematic behaviours of the actomyosin cortex in the context of the ellipsoidal shape of the embryo give rise to the orientation of an anatomical feature – the AP axis of the *C. elegans* embryo.

The explanatory power of the mathematical model arises from integrating the effects of the cortex and cytoplasm, the shape of the embryo, and the reorientation of the AP axis into a coherent whole. It demonstrated the primary importance of geometry and the action of the pseudocleavage furrow in AP axis convergence. Indeed, a minimal model based only on these two features – geometry and the furrow – recapitulates the essence of AP axis convergence. Many of the features analysed here are general to many biological systems: self organizing pattern formation (PAR domains) (Goldstein and Freeman, 1997; John Gerhart, 1997), mechanical forces (cortical flows) (Sagy et al., 2019; Etoc et al., 2016; Zhang et al., 2019; Gross et al., 2017; Münster et al., 2018), anatomy (body axes) (Hiramatsu et al., 2013; Vianello and Lutolf, 2019; Mesnard et al., 2004), and physical cues in the environment (egg geometry) (Miyazaki et al., 2015; Jülicher et al., 2018; Nitschke et al., 2020; Salbreux et al., 2022; Nitschke et al., 2022). Integration of these general features into holistic models in other experimental contexts should establish general principles of the interplay between active mechanics and geometry during development.

## Methods

*C. elegans* strains were cultured using standard culture methods (Brenner, 1974), and maintained at 20 °C. See SI: Table 1 for a list of *C. elegans* strains used in this study. RNAi interference was performed using the feeding method, see (Kamath and Ahringer, 2003). See SI: section 1 for details on RNAi experiments.

Time-lapse movies of the mid-plane cross-section of *C. elegans* embryos were obtained at room temperature. This was done using an Axio Observer Z1 - ZEISS spinning disk confocal microscope equipped with a CSU-X1 Yokogawa spinning disk head using a 63X / 1.2 NA PlanApochromat objective. PAR domains were visualized using this system equipped with an Andor iXon emCCD camera. Posteriorization of male pronucleus and cortical flows were visualized using this system equipped with a Hamamatsu ORCA-Flash4.0 V2 CMOS camera. See SI: section 1 for details on microscopy and mounting methods.

These time-lapse movies of *C. elegans* embryos were analysed to quantify the following: embryo geometry, posteriorization of the male pronucleus and cortical flows. Embryo geometry was quantified by fitting an ellipse to the boundary of the embryo mid-plane observed in the movies. Posteriorization was quantified by tracking the male pronucleus, identified as a dark circle in the cytoplasm in fluorescent microscopy images. Cortical flows were quantified similarly to (Gross et al., 2019) using particle image velocimetry on the cortical layer of the embryo, identified as a 15 pixel wide layer at the boundary of the embryo mid-plane. Only NMY-2::GFP labelled embryos were used for measuring cortical flows. See SI: section 1 for details on image analysis.

See SI: section 1 for further details on experimental methods. See SI: section 2 for details on the mathematical model and numerical simulations. See SI: section 4 for details regarding the minimal model.

## ACKNOWLEDGEMENTS

S.W.G. was supported by the DFG (grant nos. TRR 83, GR 3271/1, GR 3271/2, GR 3271/3 and GR 3271/4) and the European Research Council (grant nos. 742712 and H2020-MSCA-ITN-2015). P.G. acknowledges a EMBO Long-Term Fellowship for funding. This work was funded by the Max Planck Society and received support from the DFG under Germany’s Excellence Strategy no. EXC-2068-390729961. A.V. acknowledges support from DFG (grant no. FOR 3013). We further acknowledge computing resources provided at ZIH at TU Dresden within project WIR. We additionally thank K. Crell, T. Middelkoop, L. Pimpale, A. Narayanan and all members of the Grill Lab for experimental help and discussions. Furthermore, we thank the light microscopy facility of MPI-CBG for their support. We also thank J. White for discussions and critical inputs on the manuscript. We are grateful to P. Gönczy for comments on the manuscript.

## AUTHOR CONTRIBUTIONS

This project was conceived by A.B., M.N., P.G., M.K., M.L., A.V. and S.W.G. A.B., M.K. and P.G. designed and performed all the experiments. M.N. designed and performed all the simulations. The results were analysed by A.B., M.K., P.G. and M.N. The mathematical model was conceived by M.N., P.G., A.V. and S.W.G, while the minimal model was conceived by A.B, M.L. and S.W.G. The main text was written by A.B., M.N., M.L., A.V. and S.W.G.

